# Optimization of the glucosinolate core pathway for production of simple glucosinolates in *Escherichia coli*

**DOI:** 10.64898/2026.03.12.711178

**Authors:** Michal Poborsky, Christoph Crocoll, Barbara Ann Halkier

## Abstract

Microbial biosynthesis of plant secondary metabolites aims to provide access to compounds of medicinal or industrial value independently of the native plant. Glucosinolates are plant secondary metabolites characteristic of brassicaceous plants and recognized as promoters of human health. However, plants often contain a complex mixture of glucosinolates with insufficient amounts to elicit a clinical effect through diet. Here, we demonstrate the biosynthesis of defined glucosinolate products in *Escherichia coli* through combinatorial screening of pathway enzyme homologs, tailoring the optimal biosynthetic route for each individual product. To achieve high product titers, we establish efficient P450 expression by membrane anchor truncation and engineer sulfate assimilation to increase the supply of a sulfate donor 3’-phosphoadenosine-5’-phosphosulfate. We use benzyl glucosinolate pathway as a model to test the engineering strategies and improve the titer 37-fold over our previous study. Extrapolating the best approaches to other simple glucosinolates, we establish the first microbial synthesis of tyrosine-derived p-glucosinolate. Showing the highest titer overall, we report production of 1250 ± 91 µM indolyl-3-methyl glucosinolate, a 500-fold increase over biosynthesis in yeast.

## Introduction

Plants produce ever-growing number of recognized specialized metabolites, which, although not essential for life, provide the producer with an evolutionary advantage through their involvement in adaptation or plant defense^1^. The benefits of plant specialized metabolites extend to humans through their applications as pharmaceuticals, agrochemicals, or cosmetics. To harness their full potential, the intersection of metabolic engineering and synthetic biology aims to realize the biosynthesis of valuable plant specialized metabolites in microbial cell factories^2^. Glucosinolates are amino acid-derived, sulfur-rich specialized metabolites characteristic of brassicaceous plants. Upon consumption, glucosinolates are broken down to isothiocyanates (ITCs), which can elicit potent biological function^3^. Broccoli rich diet is associated with reduced risk of gastrointestinal cancer through epidemiological studies^4,5^ and was shown to reduce insulin resistance in type 2 diabetic patients^6^. The degradation product of tryptophan-derived indolyl-3-methyl glucosinolate (I3M) was shown to stimulate DNA damage repair in endothelial cells and act against proliferation of breast and prostate cancer^7^.

The biosynthesis of glucosinolate core structure employs eight enzymes to convert amino acids into thiohydroximates containing sulfate esters liked through the oxygen and a thioglucose on the sulfur (Figure 1)^8,9^. Carbon chain elongation of the amino acid substrate or secondary modification of the core structure further the diversity of glucosinolate products. The entrance to the pathway proceeds through two subsequent cytochrome P450 (CYP or P450) mediated steps with CYP79 enzymes converting amino acids into aldoximes, which are in turn activated into nitrile oxides by CYP83s. Exhibiting strict substrate specificity, CYP79s act as “gatekeepers” of the glucosinolate pathway, whereas the postaldoxime enzymes are not specific to the nature of the amino acid side chain^10^. However, through *Arabidopsis thaliana* co-expression analysis, the downstream enzymes are assigned to either aliphatic or aromatic (combining benzenic and indolic) glucosinolate pathways^9^. The classification system is often followed when engineering glucosinolate production in heterologous hosts^11–14^, even though it is a description of the regulation of gene expression *in planta* rather than substrate specificities of the individual enzymes.

**FIGURE 1.**
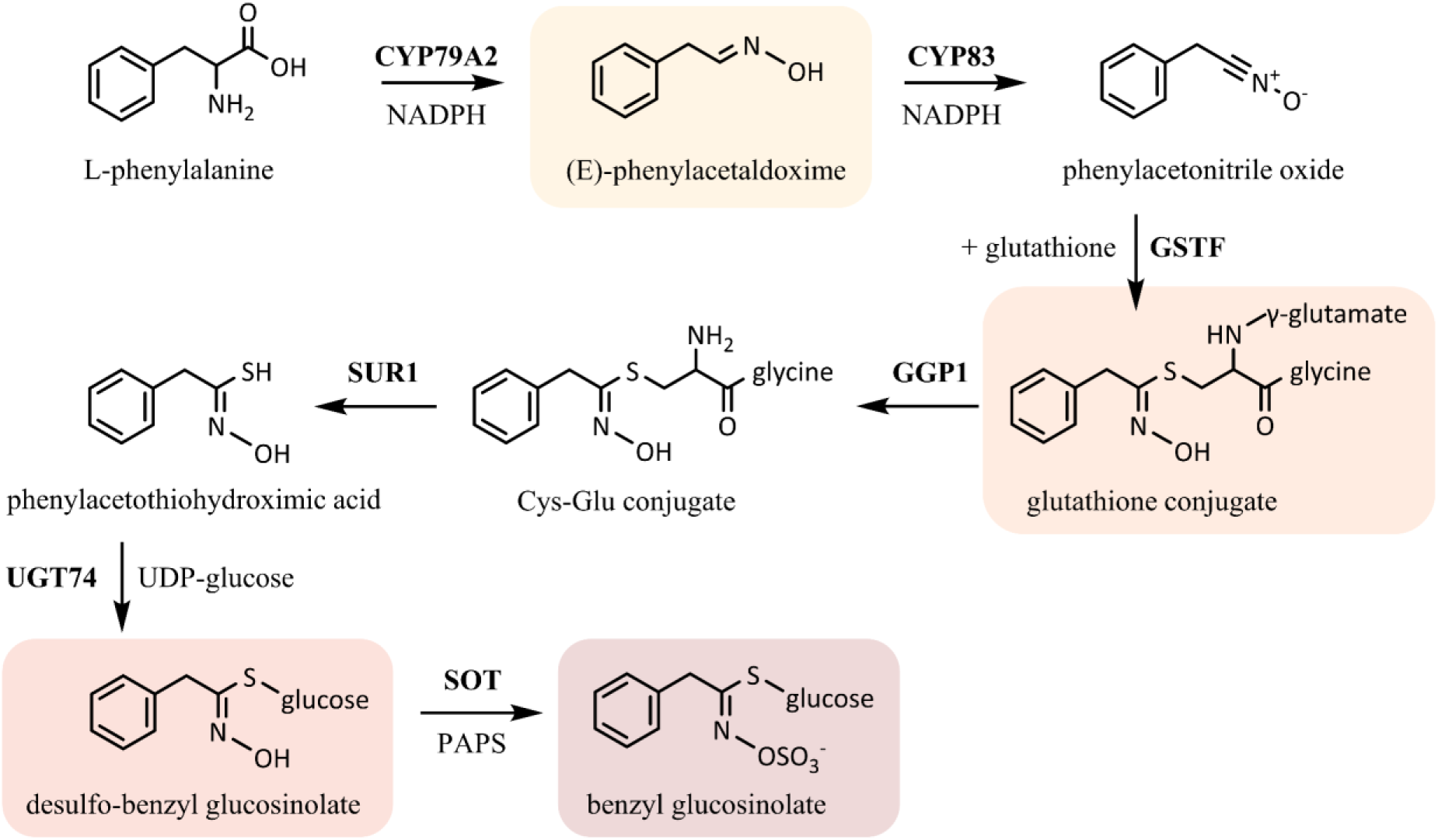
Glucosinolate core pathway with the example of phenylalanine conversion into benzyl glucosinolate. The amino acid is first converted into phenylacetaldoxime (Phe-Ox) by cytochrome P450 79A2 (CYP79A2), which is then turned into activated phenylacetonitrile oxide by another P450 from CYP83 family. In *Escherichia coli*, the functional P450s require the presence of a NADPH-dependent cytochrome P450 reductase ATR1, as there is no endogenous equivalent. Afterwards, the nitrile oxide is conjugated with glutathione into *S-*phenylacetohydroxymoyl-L-glutathione (Phe-GSH) in a reaction that can proceed spontaneously but is made more efficient when glutathione transferase (GSTF) is expressed. The glutathione moiety is first modified by gamma-glutamyl peptide 1 (GGP1) and then C-S lyase SUR1. Uridine 5’-diphospho-glucuronosyltransferase (UGT74) catalyzes addition of *S-*linked glucose, and the core structure is finalized by sulfotransferase (SOT) using PAPS as cofactor.

To date, the biosynthesis of three different glucosinolate products was reported in microbes. The tryptophan-derived I3M was produced in *Saccharomyces cerevisiae*^13^, phenylalanine-derived benzyl glucosinolate in both yeast^15^ and *E. coli*^11^ and glucoraphanin^16^, synthesized from chain-elongated methionine, in the bacterium. Achieving high productivity in microbial cell factories remains challenging as the highest titer overall was reached during benzyl glucosinolate (BGLS) production in *E. coli* at 8.3 mg/L^11^. Glucosinolate biosynthesis relies on two P450 enzymes, which are notorious for their difficult expression in yeast and *E. coli*^17,18^ and require the supply of NADPH cofactors. The glucosinolate pathway leans heavily on host resources as it incorporates glutathione, glucose moiety from uridine diphosphate glucose (UDP-glucose) and sulfate group from 3’-phosphoadenosine-5’-phosphosulfate (PAPS).

Although natively found in *E. coli* as part of sulfate assimilation, unlike in many other organisms, *E. coli* does not utilize PAPS as a sulfate donor for sulfotransferases (SOTs)^19^. Engineering of the sulfate assimilatory pathway to increase PAPS availability was shown to enhance production of sulfated compounds in the bacterium^20,21^. Even in plants, securing sufficient supply of PAPS led to 16-fold increase BGLS titers in tobacco.

Here, we describe our work towards optimization of the glucosinolate core structure pathway in *E. coli* that led to the establishment of efficient biosynthesis of three different aromatic glucosinolate products in their dedicated strains. Using BGLS pathway as a model system to evaluate our engineering strategies, we first created pathway modules to assess enzyme functionality and track intermediate progression through the pathway. By combinatorial assembly of the glucosinolate pathways with enzymes from both so-called aliphatic and aromatic pathways we identified the optimal biosynthetic route for individual products. After the optimal combination of enzymes was identified, we engineered sulfate assimilation to increase PAPS availability and applied N-terminal truncation to CYP79s and CYP83s to improve their bacterial expression. The engineered strains efficiently produced BGLS, I3M and *p-*hydroxybenzyl glucosinolate (pOHB), reaching glucosinolate titers more than 60-fold higher than previously reported. A well-functioning glucosinolate core structure pathway could pave the way for high-value glucosinolate production in microbial cell factories.

## Results

### Autoinducing medium for expression of pathway genes

In BL21 (DE3), T7 RNA polymerase expression is driven by *lacUV5* promoter and protein expression is often induced by isopropy1-β-d-thiogalactoside (IPTG). As an alternative method of induction, Studier developed autoinducing medium^22^, where the natural inducer of *lac* promoter, allolactose, drives protein expression instead. We used autoinducing media to increase the throughput of strain characterization, as it removed the need to follow culture growth and add inducer at a specific timepoint. To optimize the autoinduction protocol for P450 expression, we adjusted the tryptone and yeast extract content to match Terrific broth (TB) and tested different regimes of the growth phase. The fermentations were performed at 18°C to allow for correct folding of P450s in *E. coli*. However, as no expression should occur until late log phase with autoinducing media, we tried accelerating the cultures to a productive phase by incubating them at 28°C during the first 0, 3 and 24 h like previously reported^23^ (Figure 2). Using BGLS-producing strain to evaluate the best autoinduction protocol, any time spent at higher temperatures was detrimental to the productivity and the cells growing at 18°C during the entire 72 h fermentation outperformed the others. Incubating the cultures at 28°C for 3 hours reduced the final BGLS titer by half, and cells growing at 28°C for 24 h generated ∼90% less product. From here onwards, the fermentations were incubated at 18°C during the entire 72 h.

**FIGURE 2.**
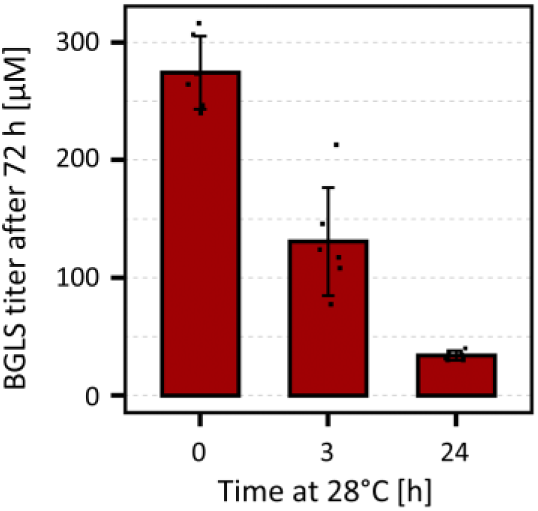
Comparison of different autoinduction protocols. Using the benzyl glucosinolate producing strain, we examined if incubating the bacteria at 28°C, 200 RPM for 3 or 24 hours at the start of the fermentation would improve product titers compared to cells maintained at 18°C during the entire 72 h.

### Identification of optimal enzyme combinations for individual glucosinolate products

For heterologous glucosinolate pathways the enzyme classification system into aromatic and aliphatic pathways was typically followed. However, Wang et al. showed that enzymes from both aromatic and aliphatic pathways can participate in the biosynthesis of BGLS as well as valine-, leucine-, and isoleucine-derived glucosinolates, and the product profile is only defined by the choice of CYP79^24^. Building on this approach, we compared a combinatorial mix of the major post-aldoxime enzymes from aliphatic and aromatic pathways when paired with CYP79A2 to produce benzyl glucosinolate and its pathway intermediates (Figure 3A).

**FIGURE 3.**
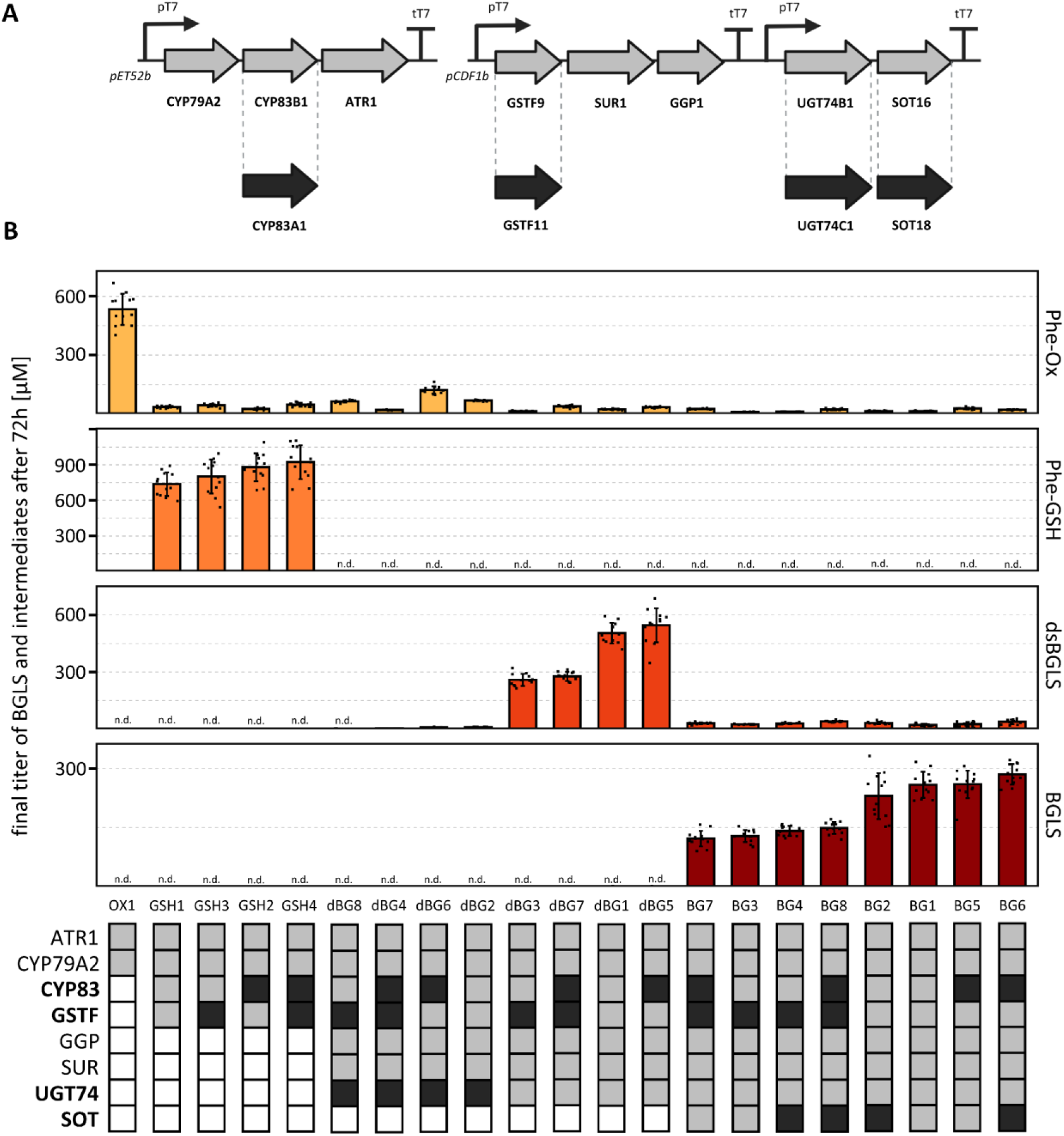
Benzyl glucosinolate (BGLS) pathway modules with a combinatorial set of aromatic and aliphatic enzymes. **A** Schematic organization of the complete BGLS pathway assembled into operons on two different plasmid backbones pET52b and pCDF1b. Variants of selected genes from so-called aromatic (light gray) and aliphatic (dark gray) pathways were used in the experiment. **B** Quantification of BGLS intermediates and products generated by BGLS pathway modules. We tracked media accumulation of phenylacetaldoxime (Phe-Ox), *S-*phenylacetohydroxymoyl-L-glutathione (Phe-GSH), desulfo-benzyl glucosinolate (dsBGLS) and benzyl glucosinolate (BGLS). Combinatorial set of selected enzymes from aromatic (light gray boxes) and aliphatic (dark gray boxes) pathways was used to assemble the modules, so functionality and substrate preferences of individual enzymes could be determined. Every strain was grown in 6 biological replicates with 2 independent experiment repeats. Outliers were defined as Q1 – 3 * IQR, Q3 + 3 * IQR (Q = quartile, IQR = interquartile range) and removed from the data. Error bars represent standard deviation from the mean.

Using phenylalanine-specific CYP79A2 and BGLS as a model pathway, we constructed different pathway modules to track accumulation of BGLS intermediates phenylacetaldoxime (Phe-Ox), *S*-phenylacetohydroxymoyl-L-glutathione (Phe-GSH), desulfo-benzyl glucosinolate (dsBGLS) as well as the final product BGLS (Figure 3B). No major bottleneck was observed as all intermediates progressed through the pathway when the necessary genes are expressed, except for dsBGLS that accumulated to 20-40 µM in strains producing BGLS. However, there was a mismatch between the maximum titers of the different pathway modules. Expressing CYP79A2 with ATR1 led to the production of 533 ± 79 µM Phe-Ox, but after the introduction of CYP83s and glutathione-S-transferase (GSTF) in the following module, Phe-GSH titers were almost twice as high. The conjugation with glutathione serves as a stabilization of the activated intermediates generated by CYP79s and CYP83s and might create a pull effect within the pathway. Conversely, in the modules downstream of Phe-GSH, the titers were steadily decreasing with a maximum dsBGLS at 546 ± 90 µM and BGLS at 285 ± 26 µM.

Comparing the enzyme homologs, we could identify the most important discriminants in the biosynthetic route to BGLS. Strains expressing UDP-glucuronosyltransferase 74C1 (UGT74C1) from the aliphatic pathway produced only trace amounts of dsBGLS or BGLS, suggesting the enzyme was not fully functional when expressed in *E. coli*. The choice of a GSTF enzyme was the second most impactful on BGLS titers, however the effect was indirect. In the Phe-GSH module, both GSTF9 and GSTF11 performed similarly, slightly in favor of GSTF11. However, when further downstream enzymes were included, the strains expressing GSTF11 produced approximately half dsBGLS and BGLS compared to GSTF9. The maximum BGLS titers were reached of 285 ± 26 µM in BG6 (Figure 3B) pathway using a combination of the aromatic enzymes with CYP83A1 and SOT18 from the aliphatic pathway.

### PAPS overaccumulation through PAPS reductase knockout

The ultimate step of the glucosinolate biosynthesis is SOT enzyme using PAPS as a sulfate donor. However, PAPS does not accumulate in *E. coli* and can be limiting for heterologous production of sulfated compounds^20,21^. Attempting to increase the PAPS availability, we disrupted the sulfate assimilatory pathway after the PAPS formation step by inactivation of PAPS reductase, generating Δ*cysH* BL21 (DE3) strain (Figure 4). However, as glucosinolate biosynthesis requires the incorporation of glutathione, a product of sulfate assimilation, we tried complementing the knockout by supplementing the fermentation medium with thiosulfate. The recently described thiosulfate sulfurtransferase (*glpE*) intersects with the sulfate pathway just below the inactivated PAPS reductase, potentially restoring any function lost in the knockout strain (Figure 4). Feeding sulfate was previously conductive to PAPS overproduction in Δ*cysH* and, therefore, it was also included in the medium.

**FIGURE 4.**
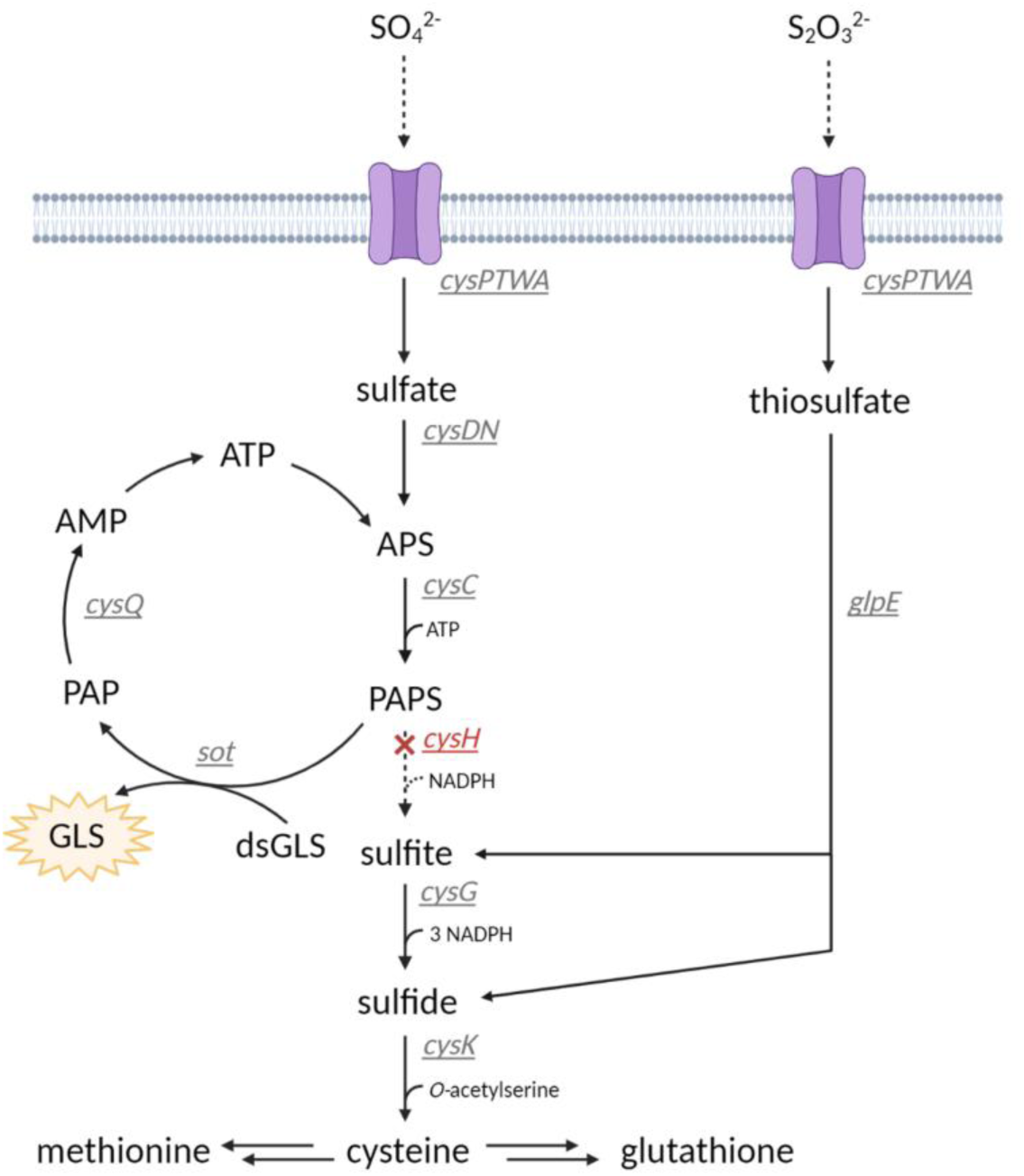
Sulfate assimilatory pathway in *E. coli* leads to the incorporation of reduced sulfur required for the amino acids cysteine and methionine and the cysteine-derived tripeptide glutathione. We generated a knockout in PAPS reductase to increase availability of PAPS for the last step of the glucosinolate core pathway, sulfurylation of desulfo-glucosinolate into intact glucosinolate. To complement the disruption of the sulfate assimilation, we fed the cells thiosulfate, which can intersect with the sulfate pathway exactly at the point of the knockout. In Δ*cysH*, each sulfur source serves a distinct function. Sulfate is only converted to PAPS and affects the conversion of dsGLS to GLS, whereas thiosulfate provides reduced sulfur for the biosynthesis of amino acids. APS = adenosine-5’-phosphosulfate, PAPS = 3’-phosphoadenosine-5’-phosphosulfate, PAP = adenosine-3’,5’-diphosphate.

As PAPS availability should directly affect conversion of dsBGLS to BGLS, we compared BGLS and dsBGLS production through the best performing BG6 pathway (Figure 3B) in wild-type and Δ*cysH* strains growing in the autoinduction medium supplemented with 5 mM SO_4_^2-^ together with or without 0.8 mM S_2_O_3_^2-^ (Table 1). The strain with inactivated PAPS reductase showed complete conversion to BGLS without thiosulfate feeding, however the pathway throughput was reduced by ∼75%, likely due to the disruption of the sulfate assimilation. Attempting to alleviate the problem by supplementing thiosulfate did not have the intended effect. Although the pathway flux was restored to original levels, 89% of the product was produced as dsBGLS, indicating problems with PAPS supply. Interestingly, with or without thiosulfate presence, the combined titers of BGLS and dsBGLS were higher in BL21 (DE3) than in Δ*cysH* strain.

**TABLE 1.** Comparison of the performance between wild-type (wt) BL21 (DE3) and Δ*cysH* strain fed with 5 mM sulfate with or without 0.8 mM thiosulfate. BGLS and dsBGLS titers were measured and the conversion rate of dsBGLS to BGLS is shown.

| Strain | SO<sub>4</sub><sup>2-</sup> [mM] | S<sub>2</sub>O<sub>3</sub><sup>2-</sup> [mM] | BGLS [μM] | dsBGLS [μM] | Conversion [%] | OD600 |
| --- | --- | --- | --- | --- | --- | --- |
| wt | 5 | 0 | 312 ± 31 | 42 ± 5 | 88.1 | 22.4 |
| cysH | 5 | 0 | 94 ± 8 | 0 | 100.0 | 19.7 |
| wt | 5 | 0.8 | 22 ± 3 | 482 ± 35 | 4.3 | 21.4 |
| cysH | 5 | 0.8 | 39 ± 8 | 324 ± 49 | 10.7 | 20.1 |

### Engineering efficient sulfate uptake and assimilation in the presence of thiosulfate

We hypothesized that thiosulfate presence in the broth interrupted the generation of PAPS from sulfate. This could happen if: (1) the sulfate/thiosulfate transporters had higher affinity towards thiosulfate, preventing sulfate entrance to the cell, (2) thiosulfate or its products downregulated the genes of sulfate assimilation. Both hypotheses are viable, as thiosulfate was shown to completely outcompete sulfate uptake in yeast^25^.

Removing any potential transcriptional regulation of sulfate assimilatory genes, we cloned *cysDNC* and *cysQ* from BL21 (DE3) genome into a synthetic operon on pCOLA-Duet plasmid (Figure 5A). *cysDNC* operon encodes genes involved in conversion of sulfate to PAPS and *cysQ* is responsible for PAPS recycling upon sulfate donation. To counteract any sulfate uptake interference, we overexpressed two different sulfate permeases (Figure 5B), *cysZ* from *Corynebacterium glutamicum* and *cysP* from *Bacillus subtillis*, where *cysZ* is not regulated by thiosulfate^26^ and *cysP* is likely not involved in thiosulfate transportation^27^. In the case thiosulfate would impact both uptake and assimilation of sulfate, we constructed pCOLA plasmids carrying *cysDNCQ* operon together with the individual sulfate permeases (Figure 5C). We engineered both wild-type and mutant BL21 (DE3).

**FIGURE 5.**
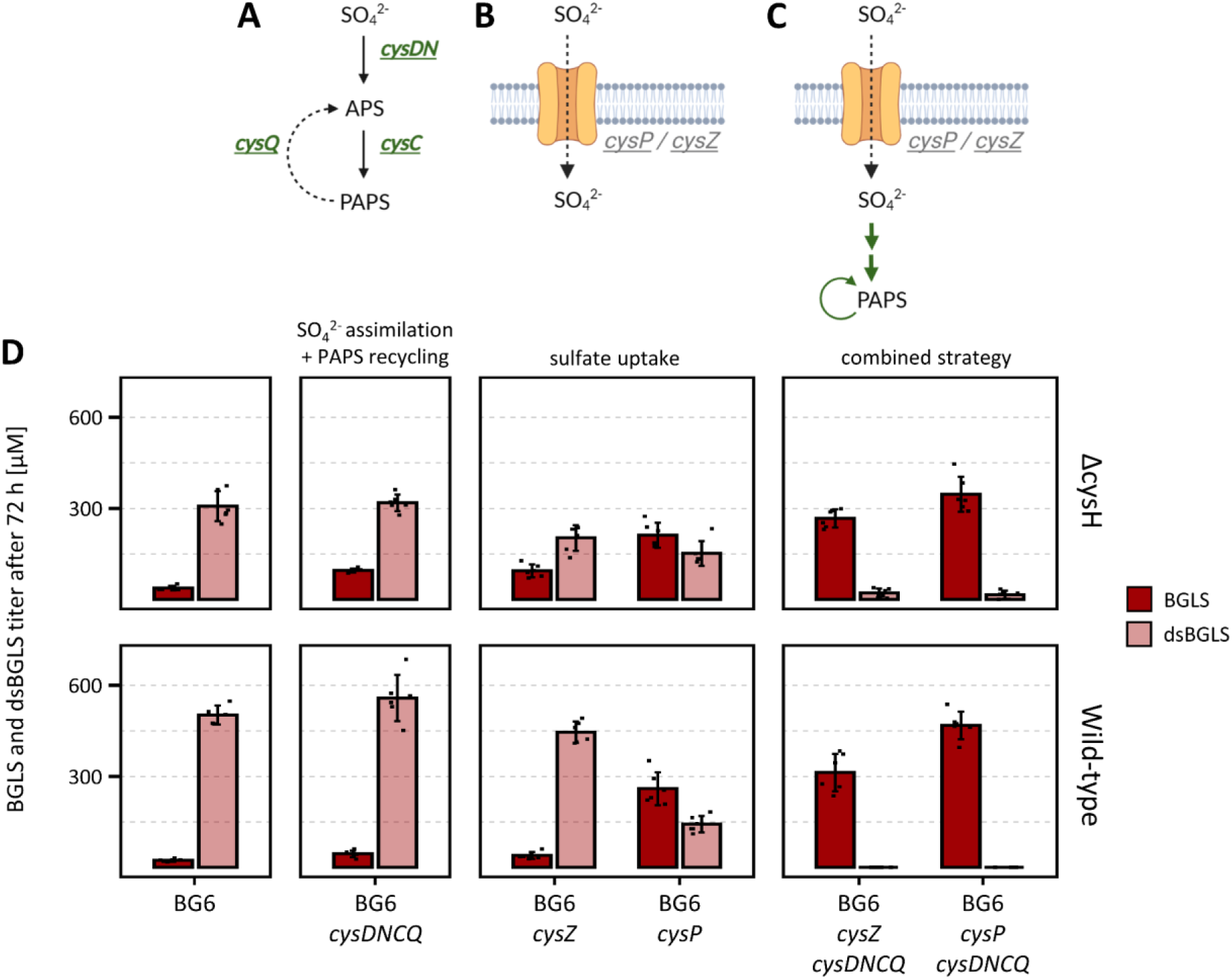
Engineering of the sulfate assimilatory pathway in wild-type and Δ*cysH* BL21 (DE3) supplemented with 5 mM sulfate and 0.8 mM thiosulfate. **A** Endogenous *cysDNC* operon was overexpressed to increase the conversion of sulfate into PAPS and to counteract any possible downregulation of the genes by thiosulfate. *cysQ* was included to initiate regeneration of PAP back to PAPS. **B** In case thiosulfate would outcompete sulfate uptake by *E. coli* transport system, two different sulfate permeases *cysZ* (*Corynebacterium glutamicum*) and *cysP* (*Bacillus subtillis*) were expressed. Both were shown to be independent of thiosulfate in their native bacteria. **C** The combination of the approaches in A and B, both sulfate uptake and assimilation were engineered. **D** Quantification of BGLS and dsBGLS in the two strains supplemented with 5 mM sulfate and 0.8 mM thiosulfate. Strains expressing BG6 pathway (Figure 3) with engineered sulfate assimilation, sulfate uptake or combination of both were compared. The bars represent mean media concentration of 6 biological replicates and the error bars represent standard deviation from the mean. APS = adenosine-5’-phosphosulfate, PAPS = 3’-phosphoadenosine-5’-phosphosulfate, PAP = adenosine-3’,5’-diphosphate.

The overexpression of *cysDNCQ* alone had a marginal impact on combined titers of dsBGLS and BGLS and only slightly improved the conversion rate in Δ*cysH* strain (Figure 5D). Comparing the two heterologous sulfate importers, *cysP* was likely more efficient in sulfate uptake as the transporter alone was able to shift the major product to BGLS. Only when simultaneously expressing sulfate assimilatory genes *cysDNC*, *cysQ* for PAPS regeneration and the sulfate permeases, the complete conversion to BGLS was achieved. The highest BGLS titer, 468 ± 45 µM, was reached by co-expressing BG6 pathway with *cysDNCQ* and *cysP* in BL21 (DE3). Improving sulfate assimilation and uptake led to a sufficient increase in PAPS availability, making the inactivation of PAPS reductase unnecessary, especially as Δ*cysH* performed worse than wild-type in every scenario.

### Establishing dedicated glucosinolate-producing strains through P450 N-terminal modification and engineering of sulfate uptake and assimilation

To demonstrate the biosynthesis of other simple glucosinolate products, we repeated the combinatorial screening of pathway enzymes, performed previously for BGLS production, with CYP79A1 and CYP79B2. CYP79A1 is specific to tyrosine and CYP79B2 only accepts tryptophan as a substrate (Table 2). The best enzyme combination for the tyrosine-derived pOHB was matching BGLS (PG6, Supplementary figure 1), whereas most I3M was produced in when GSTF11 and SOT18 were co-expressed with the enzymes from the aromatic pathway (IG4, Supplementary figure 2). Finding the best performing enzyme combination increased product titers of pOHB and I3M by ∼1.6-fold and ∼1.8-fold, respectively (Figure 6B-C). Interestingly, the optimal biosynthetic route for all aromatic glucosinolate products included some enzymes from the aliphatic pathway.

**FIGURE 6.**
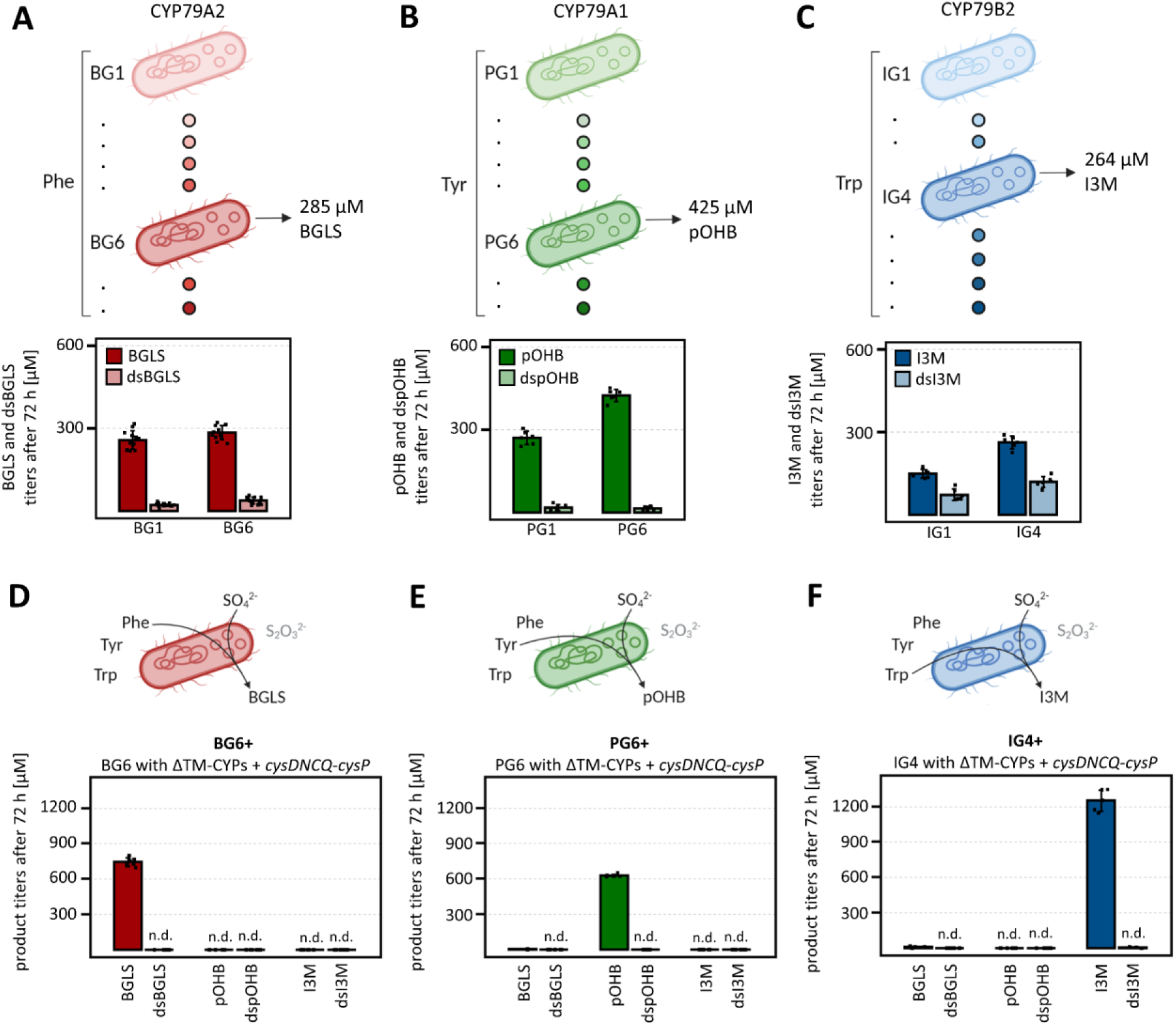
Application of the best engineering strategies to other glucosinolate products. **A, B, C** Screening of the enzyme combinations downstream of CYP79A2, CYP79A1 and CYP79B2 to identify the optimal gene set to convert phenylalanine, tyrosine, or tryptophan to simple glucosinolates. **D, E, F** Pairing of the optimal pathways for benzyl (BGLS), indolyl-3-methyl (I3M) and *p-*hydroxybenzyl (pOHB) glucosinolates with the N-terminal truncation of membrane anchors of P450 enzymes and the engineering of sulfate uptake and assimilation in BL21 (DE3) fed with 5 mM sulfate and 0.8 mM. The bars represent mean media concentration of 6 biological replicates and the error bars represent standard deviation from the mean. BG1-8 (Figure 3), PG1-8 (Supplementary figure 1) and IG1-8(Supplementary figure 2) describe different enzyme combinations in BGLS, pOHB and I3M pathways.

**TABLE 2.** Five P450 enzymes involved in simple glucosinolate pathway from three different plants were targeted for engineering in our study. Three CYP79s were selected, since they dictate which substrate can enter the pathway, whereas the subsequent CYP83s are more promiscuous. The selected range of substrates accepted by these CYP79s covers most of the proteogenic amino acid glucosinolate precursors. ΔTM represent the residues that were truncated to remove the membrane anchor.

| Enzyme | Substrates | ΔTM | Native organism | Reference |
| --- | --- | --- | --- | --- |
| CYP79A1 | L-Tyr | Δ38 | *Sorghum bicolor* | 39 |
| CYP79A2 | L-Phe | Δ14 | *Arabidopsis thaliana* | 40 |
| CYP79B2 | L-Trp | Δ42 | *Arabidopsis thaliana* | 41 |
| CYP83A1 | amino acid derived oximes | Δ23 | *Arabidopsis thaliana* | 42 |
| CYP83B1 | amino acid derived oximes | Δ21 | *Arabidopsis thaliana* | 43 |

We previously described the improvement of P450 activity by truncation of the N-terminal transmembrane helix^28^. To establish highly-productive glucosinolate cell factories, we combined all the engineering strategies together. We co-expressed the best performing pathways for BGLS, pOHB and I3M together with *cysDNCQ* and *cysP*, securing sufficient supply of PAPS by feeding sulfate and thiosulfate, and applied the N-terminal truncation to all P450s, improving their expression in *E. coli* (Figure 6D-F). Following this strategy allowed the production of 744 ± 36 µM BGLS in BG6+ strain, marking an additional ∼63% improvement compared to PAPS engineering alone. BGLS was produced as the only glucosinolate product without detectable dsBGLS accumulation. The overall engineering strategy was especially effective for I3M biosynthesis, where increasing PAPS availability and P450 truncation led to the production of 1250 ± 91 µM I3M. Compared to the original IG1 strain, the engineered IG4+ reached ∼8.4-fold higher product titer. In case of the tyrosine-derived pOHB, the improvement was less dramatic as PG6+ produced 628 ± 13 µM pOHB. BGLS, I3M and pOHB were all produced in majority in their respective strains and with no desulfo-glucosinolate accumulation. Only trace levels of BGLS were detected in IG4+ and PG6+.

## Discussion and conclusion

The promiscuity of post-aldoxime enzymes from the glucosinolate core pathway is long recognized^10^. However, when engineering glucosinolate biosynthesis in heterologous hosts, the aromatic and aliphatic categorization of the pathway enzymes was typically followed. Inspired by a recent study demonstrating that both aromatic and aliphatic pathways can accept intermediates derived from phenylalanine, leucine, isoleucine, and valine^24^, we set out to engineer glucosinolate core pathways tailored to individual products. To better understand glucosinolate biosynthesis in *E. coli*, we began by a combinatorial assembly of BGLS pathway containing post-aldoxime enzymes from aromatic and aliphatic pathways together with CYP79A2. We showed that UGT74C1, from the aliphatic pathway, was not functionally expressed in *E. coli*, consistent with a previous *in vitro* characterization of *A. thaliana* UDP-glucose-dependent glucosyltransferases^29^. Although the expression of UGT74C1 was confirmed by targeted proteomics^30^, we could only detect trace amounts of activity towards all tested substrates. As our previous attempts to realize aliphatic glucosinolate production in microbes were unsuccessful and the titers reported in the literature are very low^14,16^, we suggest the trace-level activity of UGT74C1 is the leading cause of these problems.

Curiously, the choice of a GSTF enzyme had a significant impact on BGLS production, even though GSTFs are not considered essential for glucosinolate biosynthesis, as the nitrile oxides can conjugate with glutathione spontaneously^12,13^. Even more surprisingly, GSTF9 and GSTF11 performed similarly in the module producing Phe-GSH, and the difference only appeared when additional downstream enzymes were expressed. In the Phe-GSH module, GSTF11 was slightly better than GSTF9, however, in the strains producing dsBGLS or BGLS, GSTF9 led to double product titers. In a previous study, GSTF9 expression levels were shown to correlate with CYP79A2, increasing 4-fold when the P450 was modified at the N-terminus^11^. Taken together, these results suggest that the measured effect of the GSTF enzymes is not an outcome of only their enzymatic activity or expression levels in *E. coli*, but instead a possible consequence of protein-protein interactions between GSTFs and other pathway enzymes. Many plant pathways are proposed to assemble into metabolons, large complexes of sequential metabolic enzymes, allowing the channeling of pathway intermediates^31^. Protein-protein interactions within the glucosinolate pathway were previously shown between UGT and SOT enzymes^32^, and the metabolon assembly was demonstrated in the related cyanogenic glucoside pathway^33,34^.

Although inactivation of PAPS reductase was previously used to overproduce PAPS^20^, in our system, BL21 (DE3) Δ*cysH* strain had a lower glucosinolate flux than wild-type, despite our extensive efforts. Forcing the overaccumulation of PAPS by disrupting the sulfate assimilatory pathway might have undesirable effects on the cell since inactivation of *cysH* was shown to cause cell aggregation and global changes in protein expression^19^. In contrast, the overexpression of *cysDNCQ* and *cysP* in wild-type BL21 (DE3) was a major driver behind achieving high product titers. With benefits reaching beyond a better conversion of the accumulated desulfo-glucosinolates to products, the engineered PAPS supply might create a pull effect through the glucosinolate pathway by increasing the efficiency of the ultimate sulfurylation step. Although increased glutathione availability might contribute to the better performance, our previous experiments with glutathione supplementation to the fermentation media did not show any positive effect on glucosinolate biosynthesis (data not shown).

In conclusion, we report the engineering of glucosinolate production in *E. coli* by tailoring the biosynthetic route towards three different simple glucosinolate products. We established the first microbial production of the tyrosine-derived pOHB. By engineering of PAPS availability and P450 truncation we increased the flux through the glucosinolate pathway and reached the highest glucosinolate titers in bacteria or yeast. I3M was produced at 1250 µM I3M (= 561 mg/L), more than 500-fold increase over titers from *S. cerevisiae*^13^, and high titers were reached for BGLS and pOHB at 744 µM and 628 µM, respectively.

## Methods

### Oligonucleotides, genes, plasmids, and strains

All cloning work was performed in *E. coli* NEB 10-β and BL21 (DE3) strain (both from New England BioLabs, Ipswich, USA) with pRI952 plasmid used for all fermentations. When engineering the PAPS availability, BL21 (DE3) with an inactivated PAPS reductase (*ΔcysH*) was also used. The strain was generated in a previous report^30^ by RED/ET recombination kit (Gene Bridges, Germany). pRI952 plasmid carries genes for rare tRNAs for isoleucine and arginine^35^. All plasmids and oligonucleotides used in this study can be found in Supplementary table 2 and 3, respectively. Plant genes and the gene encoding cysP sulfate permease were ordered codon optimized from either Integrated DNA Technologies (IDT, Leuven, Belgium) or Twist Bioscience (San Francisco, CA, USA). *cysDNCQ* genes were amplified from the genome of *E. coli* BL21 (DE3) and cysZ from the genome of *C. glutamicum MB001*. Oligonucleotides were purchased from IDT (Leuven, Belgium) and Sanger sequencing was provided by Eurofins Genomics (Ebersberg, Germany). Omega Bio’s bacterial kits were used for all DNA preparation. Complete overview of the overexpression targets is described in Supplementary table 1.

### DNA assembly

Proof-reading Phusion U Hot Start DNA polymerase (Thermo Fisher Scientific, Waltham, MA) with uracil containing oligonucleotides was used to amplify all DNA parts. In this study pET-52b (Novagen®, Merck, #71554), pCDF-1b (Novagen®, Merck, #71330) and pCOLADuetTM-1 (Novagen®, Merck, #71406) were used to express the genes. To assemble purified PCR products, the USER protocol was adapted from Geu Flores et al.^36^ and Cavaleiro et al.^37^ in a 10 µL reaction as follows: 1 µL USER enzyme mix , 1 µL 10x CutSmart buffer, 1 unit of DpnI, 20 ng PCR product of the plasmid backbone and 40 ng PCR product of all fragments to assemble. All component for USER assembly were purchased form New England BioLabs, Ipswich, USA. The reaction was incubated for 1 h at 37°C, then at gradually decreasing temperatures around the melting temperature (T_m_) of USER overhangs (31-26°C) with 5 min at each temperature step and finished with 30 min at 10°C. Afterwards, the entire reaction was transformed into chemically-competent *E. coli* NEB 10-β cells (New England BioLabs, Ipswich, USA). Colony PCR was performed with DreamTaq DNA polymerase (Thermo Scientific, Waltham, USA), and the sequences of cloned constructs were verified with Mix2Seq kits (Eurofins Genomics, Germany). The intergenic regions were the same in all operons to make the construction of all the pathway gene combinations straightforward. To construct the plasmids carrying two operons, the individual operons were first cloned on independent plasmids, amplified and fused together following the same USER protocol as usual. When truncating the P450 N-terminal membrane anchors, we used uniport annotations in combination with a transmembrane domain prediction tool TMHMM v2.0 (now depreciated in favor of DeepTHMHH^38^) (Table 2).

### Transformation of the expression strain

Allowing transformation of up to 4 plasmids at the same time, electrocompetent cells were always prepared fresh rather than using frozen aliquots. 30 µL of overnight culture or a single colony were inoculated into 1.4 mL LB media in a microcentrifuge tube and grown for 3-4 hours at 37°C, 900 RPM in a shaking heating block. Afterwards, the cells were washed twice with 1 mL ice-cold water and transformed with 1 µL of each plasmid. The GenePulser Xcell (Bio-Rad, Hercules, USA) electroporator was set to 1350V, 10µF, 600 Ohms. The transformants were recovered in SOC medium for 1 h at 37°C, 900 RPM, spread on LB agar with appropriate antibiotics and grown overnight at 37°C. Prior to fermentation, single colonies were picked into 1.2 mL LB media with appropriate antibiotics in a 24-well plate (Thermo Fisher Scientific, Waltham, MA) and the precultures were cultivated overnight.

### Induction of protein expression and fermentation conditions

Autoinducing medium, adapted from Studier et al.^22^, was used during all fermentations. The original recipe was adjusted to resemble Terrific broth (TB) in the yeast extract, tryptone and buffer composition, as TB is a common choice when expressing P450s. The TB-based autoinducing medium contained 24 g/L yeast extract, 12 g/L tryptone, 100 mM KH_2_PO_4_-K_2_HPO_4_ buffer pH 7.0, 0.05% glucose, 0.5% glycerol, 0.2% α-lactose, 2 mM MgSO_4_, and trace metal mix (50 µM FeCl_3_, 20 µM CaCl_2_, 10 µM MnCl_2_, 10 µM ZnSO_4_, 2 µM CoCl_2_, 2 µM CuCl_2_, 2 µM NiCl_2_, 2 µM Na_2_MoO_4_, 2 µM Na_2_SeO_3_, 2 µM H_3_BO_3_). To improve heme biosynthesis, which is necessary for functional P450 enzymes, 0.5 mM 5-aminolevulenic acid was supplied to the media. Required antibiotics were also added at the following concentrations: carbenicillin (50 µg/mL), spectinomycin (50 µg/mL), kanamycin (50 µg/mL) and chloramphenicol (34 µg/mL). Precultures were diluted 1000x into 1.2 mL autoinducing medium in round bottom 24-well plates and the fermentation was carried out in a rotary shaker at 18°C, 200 RPM for 72 h. Different autoinduction protocols were also tested to accelerate the cells towards productive phase by incubating them at 28°C, 200 RPM for the first 3 and 24 hours. Quantifying how well individual strains grew during fermentations, we measured end point OD600 after 72 h. The cultures were diluted 20-fold into 200 µL fresh media in a standard 96-well plate. When engineering the PAPS availability, potassium sulfate was supplemented in the medium at 5 mM concentration and sodium thiosulfate at 0.8 mM.

### Liquid chromatography coupled to mass spectrometry (LC-MS) metabolite analysis

#### Harvesting *E. coli* media samples

Media samples were collected after cells were spun down from 1 mL cultures in deep 96-well plates at 3700 x *g* for 5 min. The collected samples were stored in -20°C until they were analyzed by high-performance liquid chromatography tandem mass spectrometry (LC-MS/MS). Prior to analysis, an aliquot of the supernatant was diluted 25-fold in water and filtered through 0.22 µm filters.

#### Combined analysis of amino acids, benzyl glucosinolate intermediates and the three glucosinolate products by LC coupled to triple quadrupole MS (LC-MS/QqQ)

The pre-diluted media samples were diluted further in a ratio of 1:10 (v:v) in water containing the ^13^C-, ^15^N-labeled amino acid mix (Isotec, Miamisburg, US). Amino acids in the diluted extracts were directly analyzed by LC-MS/MS. The analysis method originates from a protocol described by Jander et al.^44^ and was modified from the protocol described in Docimo et al.^45^ to adjust for instrument settings and inclusion of metabolites from the benzyl-glucosinolate pathway. Chromatography was performed on an Advance ultra-high-performance liquid chromatography (UHPLC) system (Bruker, Bremen, Germany). Separation was achieved on a Zorbax Eclipse XDB-C18 column (50 x 4.6mm, 1.8µm, Agilent Technologies, Germany). Formic acid (0.05 %) in water (v/v) and acetonitrile (supplied with 0.05 % formic acid, v/v) were employed as mobile phases A and B, respectively. The elution profile was: 0-1.2 min, 3 % B; 1.2-3.8 min, 3-65 % B; 3.8-4.0 min 65-100 % B, 4.0-4.6 min 100 % B, 4.6-4.7 min 100-3 % B and 4.7-6.0 min in 3 % B. The mobile phase flow rate was 500 µl/min. The column temperature was maintained at 40°C. The liquid chromatography was coupled to an EVOQ Elite Triplequadrupole (QqQ) mass spectrometer (Bruker, Bremen, Germany) equipped with an electrospray ion source (ESI) operated in combined positive and negative ionization modes. The instrument parameters were optimized by infusion experiments with pure standards. The ion spray voltage was maintained at +3000 V or -4000V for amino acid and glucosinolate analysis, respectively. Cone temperature was set to 300°C and cone gas to 20 psi. Heated probe temperature was set to 300 °C and probe gas flow to 50 psi. Nebulizing gas was set to 60 psi and collision gas to 1.6 mTorr. Nitrogen was used as probe and nebulizing gas and argon as collision gas. Active exhaust was constantly on. Multiple reaction monitoring (MRM) was used to monitor analyte parent ion → product ion transitions: MRMs were chosen as described previously.^11,44–46^

Bruker MS Workstation software (Version 8.2.1, Bruker, Bremen, Germany) was used for data acquisition and processing. Linearity in ionization efficiencies were verified by analyzing dilution series of standard mixtures (amino acid standard mix, Fluka plus Gln and Trp, also Fluka). All samples were spiked with ^13^C-, ^15^N-labeled amino acids at a concentration of 10 µg/mL. The concentration of the individual labeled amino acids in the mix had been determined by comparison to a reference standard by LC-MS/MS analysis. Individual amino acids in the sample were quantified by the respective ^13^C-, ^15^N-labeled amino acid internal standards, except for tryptophan and asparagine: tryptophan was quantified using ^13^C-, ^15^N-Phe applying a response factor of 0.42, asparagine was quantified using ^13^C-, ^15^N-Asp applying a response factor of 1.0. Additional MRMs for glucosinolate intermediates like aldoximes and GSH conjugates were included into the method.

