## Supplementary Figures and Tables for "Optimization of the glucosinolate core pathway for production of simple glucosinolates in *Escherichia coli*"

### Supplementary materials

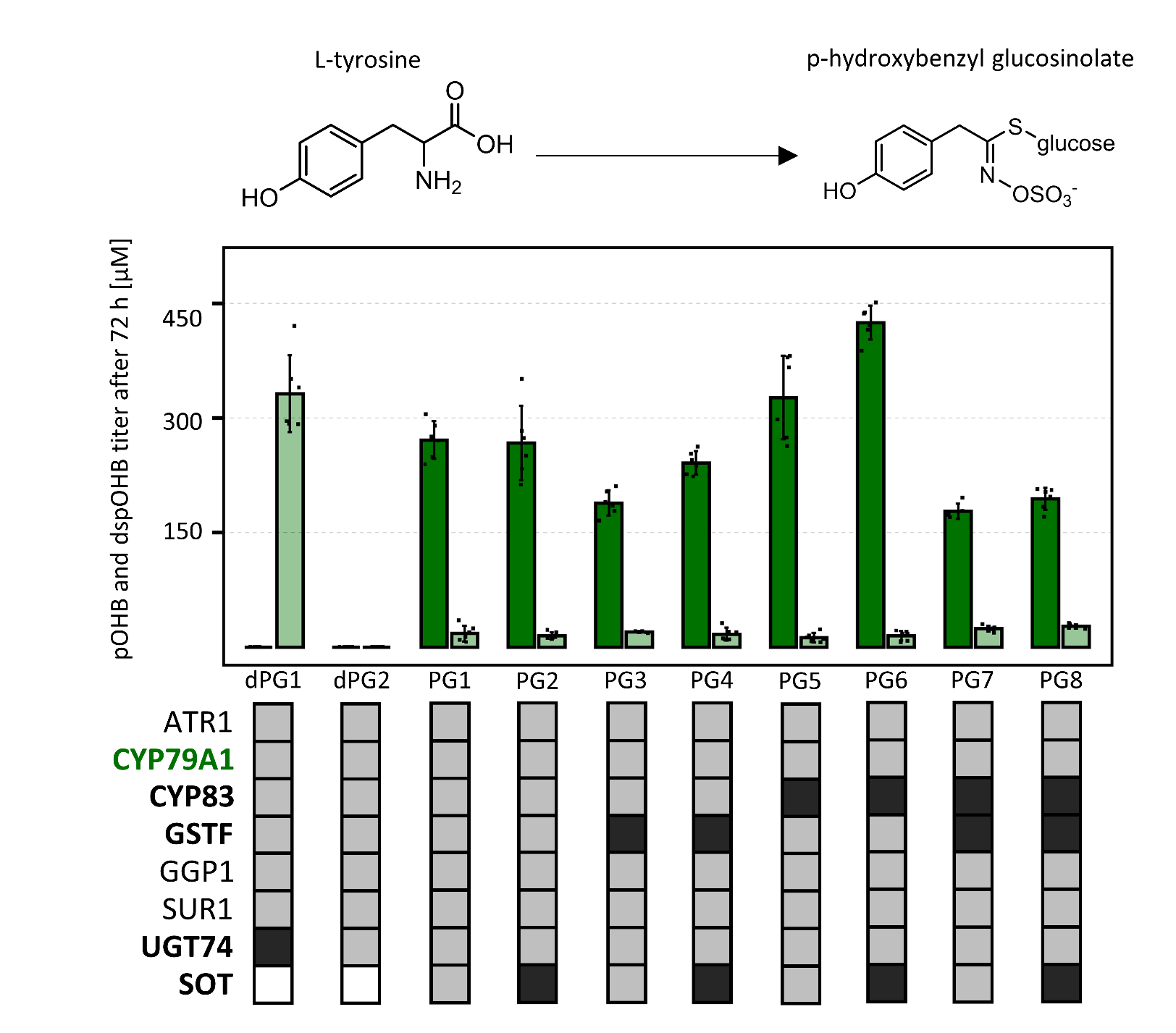

**Supplementary figure 1 |** Combinatorial screening of glucosinolate core structure pathway enzymes downstream of CYP79A1 for the production of tyrosine-derived *p-*hydroxybenzyl glucosinolate. The operons were organized in the same way as in Figure 3A, but CYP79A2 was substituted with CYP79A1. Combinatorial set of selected enzymes from aromatic (light gray boxes) and aliphatic (dark gray boxes) pathways was used to assemble the pathways. Every strain was grown in 6 biological replicates. The bars represent mean concertation values and the error bars standard deviation from the mean.

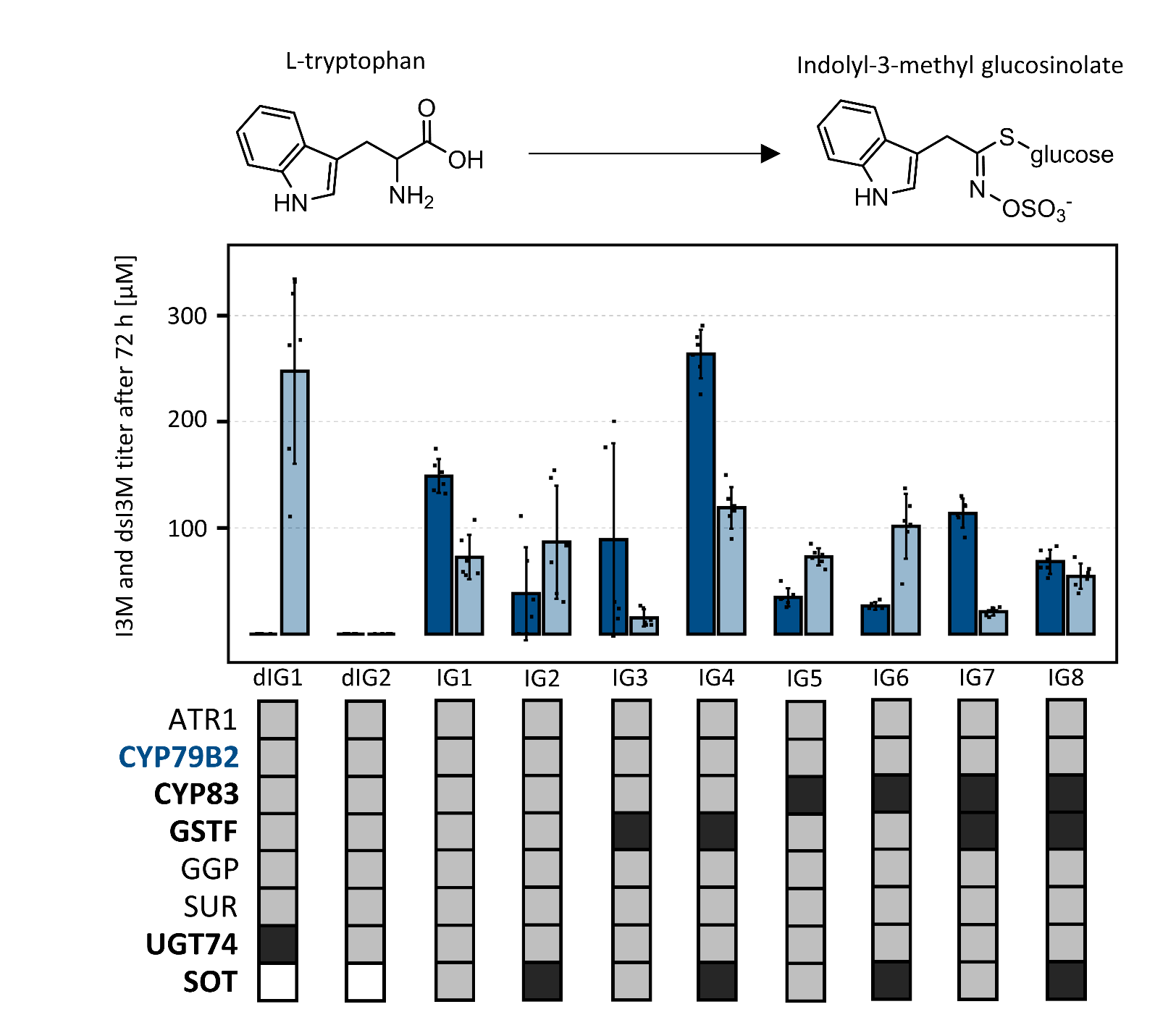

**Supplementary figure 2 |** Combinatorial screening of glucosinolate core structure pathway enzymes downstream of CYP79B2 for the production of tryptophan-derived indolyl-3-methyl glucosinolate. The operons were organized in the same way as in Figure 3A, but CYP79A2 was substituted with CYP79B2. Combinatorial set of selected enzymes from aromatic (light gray boxes) and aliphatic (dark gray boxes) pathways was used to assemble the pathways. Every strain was grown in 6 biological replicates. The bars represent mean concertation values and the error bars standard deviation from the mean.

**SUPPLEMENTARY TABLE 1 |** List of overexpressed proteins in the study.

| **Gene** | **Protein** | **Organism** | **Uniprot ID** | **DNA source** |
| --- | --- | --- | --- | --- |
| CYP79A2 | Phenylalanine N-monooxygenase | *Arabidopsis thaliana* | Q9FLC8 | codon optimized from Integrated DNA Technologies |
| CYP79A1 | Tyrosine N-monooxygenase | *Sorghum bicolor* | Q43135 | codon optimized from Twist Bioscience |
| CYP79B2 | Tryptophan N-monooxygenase | *Arabidopsis thaliana* | O81346 | codon optimized from Twist Bioscience |
| CYP79D2 | Valine N-monooxygenase 2 | *Manihot esculenta* | Q9M7B7 | codon optimized from Twist Bioscience |
| CYP83B1 | Cytochrome P450 83B1 | *Arabidopsis thaliana* | O65782 | codon optimized from Integrated DNA Technologies |
| CYP83A1 | Cytochrome P450 83A1 | *Arabidopsis thaliana* | P48421 | codon optimized from Integrated DNA Technologies |
| GSTF9 | Glutathione S-transferase F9 | *Arabidopsis thaliana* | O80852 | codon optimized from Integrated DNA Technologies |
| GSTF11 | Glutathione S-transferase F11 | *Arabidopsis thaliana* | Q96324 | codon optimized from Integrated DNA Technologies |
| GGP1 | Gamma-glutamyl peptidase 1 | *Arabidopsis thaliana* | Q9M0A7 | codon optimized from Integrated DNA Technologies |
| SUR1 | S-alkyl-thiohydroximate lyase 1 | *Arabidopsis thaliana* | Q9SIV0 | codon optimized from Integrated DNA Technologies |
| UGT74B1 | UDP-glycosyltransferase 74B1 | *Arabidopsis thaliana* | O48676 | codon optimized from Integrated DNA Technologies |
| UGT74C1 | UDP-glycosyltransferase 7CB1 | *Arabidopsis thaliana* | Q9SKC1 | codon optimized from Integrated DNA Technologies |
| SOT16 | Cytosolic sulfotransferase 16 | *Arabidopsis thaliana* | Q9C9D0 | codon optimized from Integrated DNA Technologies |
| SOT18 | Cytosolic sulfotransferase 18 | *Arabidopsis thaliana* | Q9C9C9 | codon optimized from Integrated DNA Technologies |
| cysN | Sulfate adenylyltransferase subunit 1 | *Escherichia coli* BL21 (DE3) | P23845 | amplified from genome of BL21 strain |
| cysD | Sulfate adenylyltransferase subunit 2 | *Escherichia coli* BL21 (DE3) | P21156 | amplified from genome of BL21 strain |
| cysC | Adenylyl-sulfate kinase | *Escherichia coli* BL21 (DE3) | P0A6J1 | amplified from genome of BL21 strain |
| cysQ | 3'(2'),5'-bisphosphate nucleotidase | *Escherichia coli* BL21 (DE3) | P22255 | amplified from genome of BL21 strain |
| cysP | Sulfate permease | *Bacillus subtilis* | O34734 | codon optimized from Twist Bioscience |
| **Gene** | **Protein** | **Organism** | **Locus** | **DNA source** |
| cysZ | Putative sulphate permease | *Corynebacterium glutamicum* | CGP_RS12940 | MB001 strain genome |

**SUPPLEMENTARY TABLE 2 |** List of all plasmids created and used during this work.

| **Plasmid** | **Reference** |
| --- | --- |
| pCDF1b | Novagen |
| pET52b(+) | Novagen |
| pCOLA-Duet1b | Novagen |
| pRI952 | Del Tito et al., 1995 |
| pET52-CYP79A2-ATR1 | This study |
| pET52-CYP79A2-CYP83B1-ATR1 | This study |
| pET52-CYP79A2-CYP83A1-ATR1 | This study |
| pET52-SbCYP79A1-CYP83B1-ATR1 | This study |
| pET52-SbCYP79A1-CYP83A1-ATR1 | This study |
| pET52-CYP79B2-CYP83B1-ATR1 | This study |
| pET52-CYP79B2-CYP83A1-ATR1 | This study |
| pET52-CYP79D2-CYP83B1-ATR1 | This study |
| pET52-CYP79D2-CYP83A1-ATR1 | This study |
| pET52-Δ14CYP79A2-Δ23CYP83B1-ATR1 | This study |
| pET52-Δ14CYP79A2-Δ21CYP83A1-ATR1 | This study |
| pET52-Δ38SbCYP79A1-Δ23CYP83B1-ATR1 | This study |
| pET52-Δ38SbCYP79A1-Δ21CYP83A1-ATR1 | This study |
| pET52-Δ42CYP79B2-Δ23CYP83B1-ATR1 | This study |
| pET52-Δ42CYP79B2-Δ21CYP83A1-ATR1 | This study |
| pET52-Δ41CYP79D2-Δ23CYP83B1-ATR1 | This study |
| pET52-Δ41CYP79D2-Δ21CYP83A1-ATR1 | This study |
| pCDF-GSTF9 | This study |
| pCDF-GSTF11 | This study |
| pCDF-[GSTF9-SUR1-rGGP1]-[UGT74B1] | This study |
| pCDF-[GSTF9-SUR1-rGGP1]-[UGT74B1-SOT16] | This study |
| pCDF-[GSTF9-SUR1-rGGP1]-[UGT74B1-SOT18] | This study |
| pCDF-[GSTF9-SUR1-rGGP1]-[UGT74C1] | This study |
| pCDF-[GSTF11-SUR1-rGGP1]-[UGT74B1] | This study |
| pCDF-[GSTF11-SUR1-rGGP1]-[UGT74B1-SOT16] | This study |
| pCDF-[GSTF11-SUR1-rGGP1]-[UGT74B1-SOT18] | This study |
| pCDF-[GSTF11-SUR1-rGGP1]-[UGT74C1] | This study |
| pCOLA-cysDNCQ | This study |
| pCOLA-cysZ[C.glut] | This study |
| pCOLA-cysP[B.subt] | This study |
| pCOLA-cysDNCQ-cysZ | This study |
| pCOLA-cysDNCQ-cysP | This study |

**SUPPLEMENTARY TABLE 3 |** List of all oligonucleotides used in the study.

| **Oligonucleotide** | **Sequence** |
| --- | --- |
| USERA1-pET52-rev | atctccttctuaaagttaaacaaaattatttctagaggggaattg |
| USERB-pET52-fwd | acatgtgttcugctgccaccgctgag |
| USERC-pET52-fwd | acttgttcagugctgccaccgctgag |
| USERD-pET52-fwd | attggactctugctgccaccgctgag |
| USERA2-pCDF-rev | atatctccttautaaagttaaacaaaattatttctacagggg |
| USERB-pCDF-fwd | actgttgataucgctgagcaataactagcataacc |
| USERC-pCDF-2-fwd | aagaagtcatucgctgagcaataactag |
| USERD-pCDF-3-rev | atctaactcatucgctgagcaataactagc |
| USERA2-pCOLA-rev | atatctccttautaaagttaaacaaaattatttctacagggg |
| USERB-pCOLA-fwd | acctaatgtcuatgctgccaccgctgag |
| USERC-pCOLA-2-fwd | agtgaattccuatgctgccaccgctg |
| USERA1-CYP79A2-fwd | aagaaggagauatacatatgttggcctttattattggactg |
| USERA1-Δ14CYP79A2-fwd | aagaaggagauatacatatgaaacgcaaagaaaaaaaaaagacg |
| USERB-CYP79A2-rev | agaacacatguttaggtcggataaacgtgc |
| USERA1-SbCYP79A1-fwd | aagaaggagauatacatatggcaaccatggaagtagaagccg |
| USERA1-Δ38SbCYP79A1-fwd | aagaaggagauatacatatgcgcgcgttacgtcgcc |
| USERB-SbCYP79A1-rev | agaacacatguttagatgctaatgcttgggtacagatggg |
| USERA1-CYP79B2-fwd | aagaaggagauatacatatgaacacgtttactagtaattcg |
| USERA1-Δ42CYP79B2-fwd | aagaaggagauatacatatgaagaagttaatgacggacc |
| USERB-CYP79B2-rev | agaacacatgutcacttaaccgtggggtataaatgc |
| USERA1-CYP79D2-fwd | aagaaggagauatacatatggctatgaacgttagtaccaccgc |
| USERA1-Δ41CYP79D2-fwd | aagaaggagauatacatatgaaattgcagaagcgtgc |
| USERB-CYP79D2-rev | agaacacatguttacggagacgttggatataaatgcgg |
| USERB-CYP83B1-fwd | acatgtgttcuttaagaaggagatataccatggacctcttactgattattgc |
| USERB-d23CYP83B1-fwd | acatgtgttcuttaagaaggagatataccatgaaaaagtccttacgtttacc |
| USERC-CYP83B1-rev | actgaacaagutcaaatatgtttggttggtgccag |
| USERB-CYP83A1-fwd | acatgtgttcuttaagaaggagatataccatggaggacattatcattgggg |
| USERB-d21CYP83A1-fwd | acatgtgttcuttaagaaggagatataccatgaaaccgaaaaccaaacgc |
| USERC-CYP83A1-rev | actgaacaaguttaatacttgtttaccttctctggcacc |
| USERA2-GSTF9-fwd | ataaggagatauacatatggttctgaaggtttacgg |
| USERB-GSTF9-rev | agaacacatguttacgccgggaacgag |
| USERA2-GSTF11-fwd | ataaggagatauacatatggtggtgaaggtctatgg |
| USERB-GSTF11-rev | agaacacatguttagtaagcggccagttcc |
| USERC-rGGP1-fwd | acttgttcaguttaagaaggagatataccatggtggagcaaaagcg |
| USERD-rGGP1-rev | aagagtccaautcaattcgtcggcacacgac |
| USERB-SUR1-fwd | acatgtgttcuttaagaaggagatataccatgagcgaagaacaac |
| USERC-SUR1-rev | actgaacaaguttacatttccaggttattgtccg |
| USERA2-UGT74B1-fwd | ataaggagatauactatggcagaaaccacgccg |
| USERB-UGT74B1-rev | agaacacatguttatttgcccagactctcaatgaattcg |
| USERA2-UGT74C1-fwd | ataaggagatauactatgagcgaggcgaagaaagg |
| USERB-UGT74C1-rev | agaacacatguttaggtaagcagcgctacgaac |
| USERB-SOT16-fwd | acatgtgttcuttagaaggagatatacatatggaaagcaaaacgaccc |
| USERC-SOT16-rev | actgaacaaguttaattatcatgctgcagcaacccg |
| USERB-SOT18-fwd | acatgtgttcuttagaaggagatatacatatggaaagcgagaccttaacg |
| USERC-SOT18-rev | actgaacaaguttatttgccgtgttcgagcaggc |
| USERA2-cysDNC-fwd | ataaggagatauacatatggatcaaatacgacttactcacc |
| USERB-cysDNC-rev | agaacacatgutcaggatctgataatatcgttctgtc |
| USERB-cysQ-fwd | acatgtgttcuaagttaacaccgctcacag |
| USERC-cysQ-rev | actgaacaaguttaataaatagacactctgaatcccg |
| USERA2-cysZ-fwd | ataaggagatauacatatgcagacattaatctttatcgcc |
| USERB-cysZ-rev | agaacacatguttaagtgcttttaatcttttggccc |
| USERA2-cysP-fwd | ataaggagatauacatatggaattagccgctattttatttagc |
| USERB-cysP-rev | agaacacatgutcagattcccccgcc |

s
